# Species Designations Belie Phenotypic and Genotypic Heterogeneity in Oral Streptococci

**DOI:** 10.1101/375139

**Authors:** Irina M. Velsko, Brinta Chakraborty, Marcelle M. Nascimento, Robert A. Burne, Vincent P. Richards

## Abstract

Health-associated oral *Streptococcus* species are promising probiotic candidates to protect against dental caries. Ammonia production through the arginine deiminase system (ADS), which can increase the pH of oral biofilms, and direct antagonism of caries-associated bacterial species are desirable properties for oral probiotic strains. ADS and antagonistic activities can vary dramatically among individuals, but the genetic basis for these differences is unknown. We sequenced whole genomes of a diverse set of clinical oral *Streptococcus* isolates and examined the genetic basis of variability in ADS and antagonistic activities. A total of 113 isolates were included and represented ten species: *S. australis*, A12-like, *S. cristatus*, *S. gordonii*, *S. intermedius*, *S. mitis*, *S. oralis* including *S. oralis* subsp. *dentisani*, *S. parasanguinis, S. salivarius*, and *S. sanguinis*. Mean ADS activity and antagonism on *Streptococcus mutans* UA159 were measured for each isolate, and each isolate was whole genome-shotgun sequenced on an Illumina MiSeq. Phylogenies were built of genes known to be involved in ADS activity and antagonism. Several approaches to correlate the pan-genome with phenotypes were performed. Phylogenies of genes previously identified in ADS activity and antagonism grouped isolates by species but not by phenotype. GWAS identified additional genes potentially involved in ADS activity or antagonism across all the isolates we sequenced as well as within several species. Phenotypic heterogeneity in oral streptococci is not necessarily reflected by genotype and is not species-specific. Probiotic strains must be carefully selected based on characterization of each strain, and not based on inclusion within a certain species.

**Importance:** Representative type strains are commonly used to characterize bacterial species, yet species are phenotypically and genotypically heterogeneous. Conclusions about strain physiology and activity based on a single strain therefore may be inappropriate and misleading. When selecting strains for probiotic use, the assumption that all strains within a species share the same desired probiotic characteristics share those characteristics may result in selection of a strain that lacks the desired traits, and therefore makes a minimally effective or ineffective probiotic. Health-associated oral streptococci are promising candidates for anti-caries probiotics, but strains need to be carefully selected based on observed phenotypes. We characterized the genotype and anti-caries phenotypes of strains from ten species of oral Streptococci and demonstrate poor correlation between genotype and phenotype across all species.

## Introduction

Dental caries is a significant health problem and the most common oral infectious disease, causing substantial morbidity worldwide. Caries develop when the tooth enamel is demineralized through successive exposure to low pH, a condition driven by fermentation of dietary carbohydrates into organic acids by acidogenic oral bacterial species. Treatment for caries can be expensive, and disease prevention is a major goal of oral healthcare research.

The use of orally-administered probiotic species is gaining popularity as a strategy for maintaining oral health. This involves introducing bacterial strains to the oral cavity with the goal of promoting growth and metabolic activity of a health-associated oral biofilm, while suppressing growth and metabolic activity of disease-associated species. Several studies have demonstrated successful *in vivo* and *in vitro* application of dairy product-derived oral probiotic species, predominantly lactobacilli, highlighting the potential of probiotics in oral healthcare. Select strains of lactobacilli inhibit growth and biofilm formation of caries-associated species *Streptococcus mutans* and *Candida albicans* in culture (1-3), which is a prime caries prevention strategy. *In vitro* biofilm growth assays demonstrated that strains of *Lactobacillus*, *Lactococcus*, and *Streptococcus* can integrate into saliva-derived or defined-species biofilms and are maintained in the biofilms over several days (4-8). However, an *in vivo* study reported that no probiotic lactobacilli were detected in dental plaque of individuals after 8-day treatment with fermented milk (9), so the method by which such probiotic strains act on the biofilm *in vivo* needs to be further investigated.

In addition to food-derived probiotic strains, there are many bacterial species in dental plaque that are associated with health, which may be mined for probiotic potential. These oral species have the advantage of being adapted to growth in the mouth and the oral biofilm, and may offer more sustainable and longer-term probiotic benefits than species from external sources like dairy products. In particular, several *Streptococcus* species including *S. gordonii*, *S. sanguinis*, and *S. salivarius* are associated with oral health (10-12), and *S. salivarious* K12 has been adapted as a probiotic for pharyngitis/tonsillitis (13), halitosis (14), and otitis media (15).

Buffering biofilm pH through ammonia production is a promising health-associated activity of oral Streptococci (10). The arginine deiminase system (ADS) is a dominant method used by Streptococci to produce ammonia from arginine. This pathway has been extensively characterized in *S. gordonii* (16-19) and consists of an operon containing five genes encoding structural proteins: *arcA* (arginine deiminase), *arcB* (ornithine carbamoyltransferase), *arcC* (carbamate kinase), *arcD* (arginine-ornithine antiporter), and *arcT* (putative transaminase or peptidase); and two regulatory genes immediately downstream that are co-transcribed in the opposite direction from the operon: *arcR* and *queA*, which are essential for optimal ADS activity in *S. gordonii* (16). Additionally, upstream of *arcA* is *flp*, another regulatory element involved in ADS activity (17). Expression of the ADS operon is regulated by environmental factors, including the presence of arginine (18), sugar carbon source (17), and the presence of oxygen (18, 20). Strains with defective ADS expression or regulation are more sensitive to pH-induced killing (19), suggesting this pathway is important for maintaining health-associated species in the presence of acidogenic species.

Use of arginine-containing toothpaste and mint prebiotics to boost ADS activity in plaque and protect against caries development and progression is also a promising method to promote health-associated activity in oral biofilms (21-24). As arginine directly affects growth and pathogenesis of *S. mutans* (25), developing probiotics that target arginine metabolism maybe especially effective in preventing caries (26). Arginine catabolism is clinically relevant to caries development. Clinical studies have shown that ADS activity is higher in plaque and saliva of patients who have never had caries than patients with active caries, both in adults (27) and children (28). Further, Nascimento, *et al.* (2009) found an inverse relationship between ADS activity and abundance of *S. mutans* in plaque samples. Unexpectedly, they found no correlations between the abundance of health-associated *Streptococcus* species and ADS activity level, while some plaque samples from caries sites had high ADS activity (28). They concluded that there must be more to ADS activity than simply the presence or abundance of health-associated streptococci.

Recent phenotypic characterization of ADS activity in a variety of oral *Streptococcus* species grown in different conditions (arginine availability, pH, carbohydrate source, oxygen tension) showed substantial variation in activity within and between species and within growth conditions (29). This confirmed the conclusion of Nascimento, *et al*. (2013) that no health-associated *Streptococcus* species are collectively associated with ADS activity, rather, ADS activity is highly strain-specific. In addition, these clinical isolates had a range of ability to antagonize growth of *S. mutans* (29), also a desirable trait in a probiotic strain. The genetic basis for variability in ADS activity and *S. mutans* antagonism has not yet been examined, but may provide insight for probiotic development. Rational selection of probiotic strains is particularly important because of the genotypic and phenotypic heterogeneity within oral *Streptococcus* species. Here we examined the probiotic properties and genome composition of a wide variety of oral *Streptococcus* species isolated from dental plaque. We show substantial phenotypic and genotypic heterogeneity of all species examined, which has implications for targeted probiotic strain selection.

## Results

### Species assignments

A total of 113 *Streptococcus* species were isolated from supragingival dental plaque samples, characterized for ADS activity and antagonism on and by *S. mutans* UA159, and whole genome shotgun sequenced. Nine species were identified by 16S rRNA gene sequencing of 106 isolates, and seven isolates could not be identified at species-level. Core genome analysis confirmed that we characterized and sequenced two *S. australis*, two A-12-like isolates, eleven *S. cristatus*, seventeen *S. gordonii*, eleven *S. intermedius*, twenty-seven *S. mitis*, eight *S. oralis*, six *S. oralis* subsp. *dentisani*, twenty-five *S. sanguinis*, one each of *S. parasanguinis* and *S. salivarius*, and two for which a species could not be identified as they grouped with the *S. mitis/S. oralis* complex in the phylogeny (Figure S1, Table S1). Given that previous work has placed *S. oralis* subsp. *dentisani* as a distinct subclade of *S. oralis* (50), we performed all analyses by both grouping all together and keeping them as separate groups. Likewise, the *S. australis* and A12-like isolates were grouped together for all analyses because we had only two isolates of each and they more closely related to each other than to other *Streptococcus* species (51). Based on the core genome phylogeny of our isolates, the phylogenetic relationships of the species we sequenced follow the branching patterns reported for these species within the genus *Streptococcus* (52) with the exception that *S. oralis*, *S. mitis*, and *S. oralis* subsp. *dentisani* are intermixed within their clade with no clear species groupings (Figure S1).

### *Heterogeneity of arginine deiminase activity and antagonism on* S. mutans *within diverse* Streptococcus *species*

Phenotypic heterogeneity was shown by the range of ADS activity and antagonism on *S. mutans* antagonism within strains of each species (Figure 1, Table 1). As arginine deiminase activity was first described in *S. gordonii* DL1, we used *S. gordonii* as the reference group for our statistical tests. *Streptococcus mitis*, *S. oralis*, and *S. oralis* + *S. oralis* subsp. *dentisani* each had significantly lower ADS activity by one-way ANOVA than *S. gordonii* (Figure 1A, Table 1). The large standard deviations demonstrate substantial species phenotypic diversity, particularly in *S. gordonii* and *S. sanguinis*. A single isolate of *S. oralis* subsp. *dentisani* had exceptionally high ADS activity for the species (half-filled circle Figure 1), and was responsible for that group’s large standard deviation. The *S. australis* and A12-like isolates separated into two clusters, with the A12-like isolates (half-filled circles Figure 1A) exhibiting higher average ADS activity than the *S. australis* isolates (filled circles).

**Figure 1.**
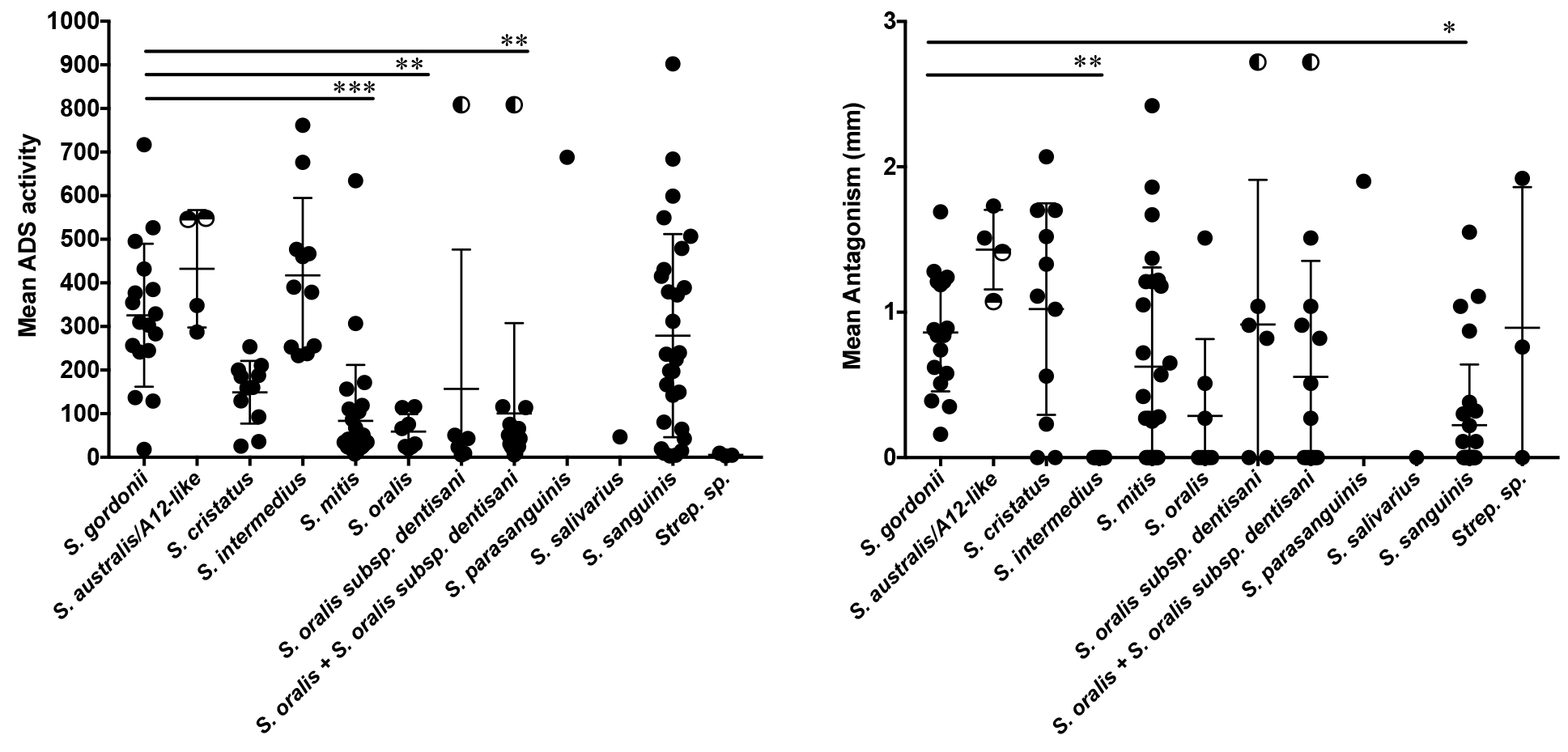
Phenotypic diversity within diverse clinical oral streptococcus isolates. A. Mean ADS activity of each isolate included in this study. B. Mean antagonism of *S. mutans* UA159 of each isolate included in this study. Half-filled circles in *S. australis*/A12-like indicate the A12-like isolates. Half-filled circles in *S. oralis* subsp. *dentisani* and *S. oralis* + *S. oralis* subsp. *dentisani* indicate the same isolate. * p < 0.05, ** p < 0.01, *** p < 0.001.

**Table 1.**
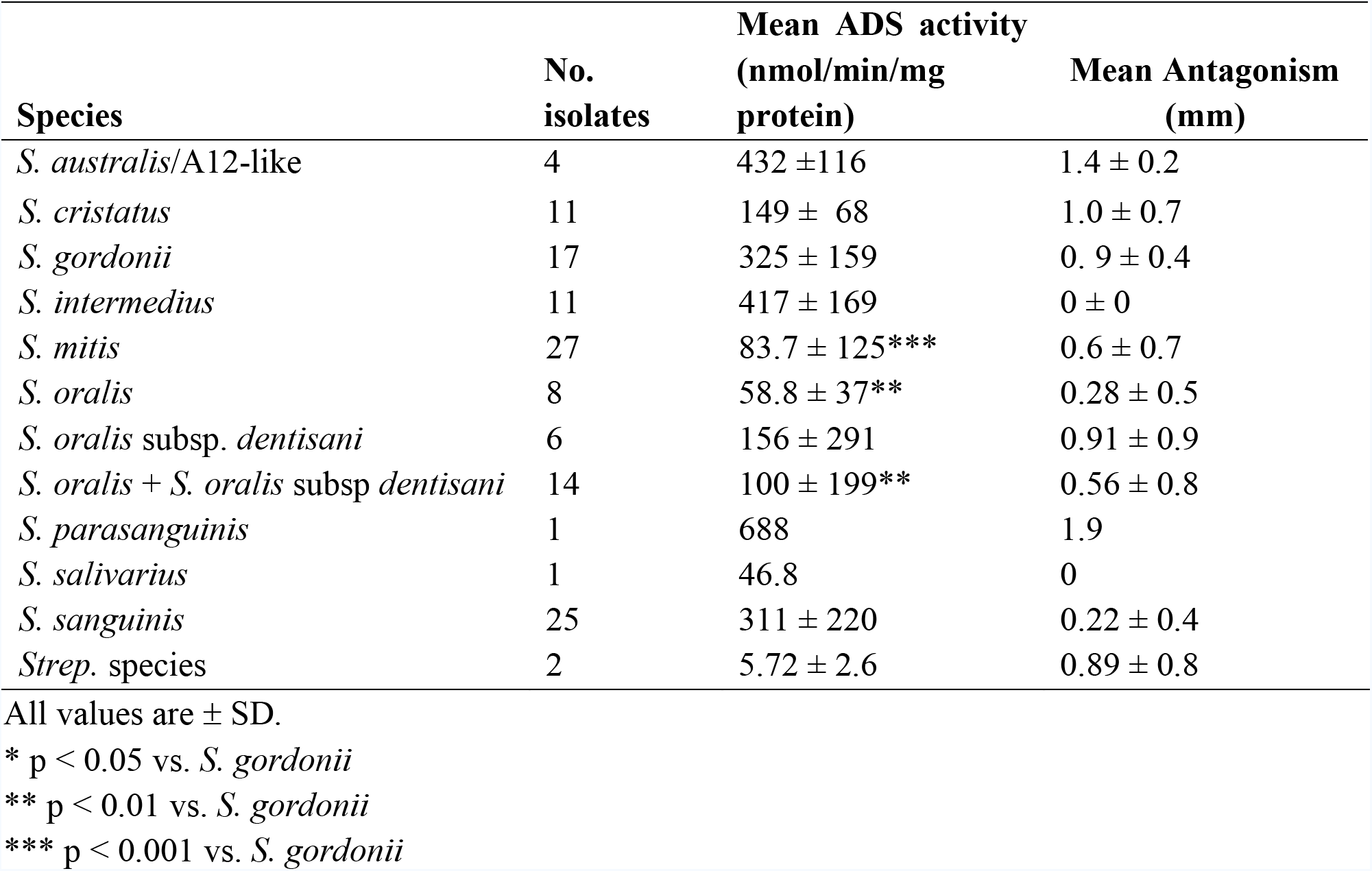
Mean arginine deiminase activity and *S. mutans* antagonism of diverse *Streptococcus* species. All values are ± SD. * p < 0.05 vs. *S. gordonii* ** p < 0.01 vs. *S. gordonii* *** p < 0.001 vs. *S. gordonii*

| Species | No. isolates | Mean ADS activity (nmol/min/mg protein) | Mean Antagonism (mm) |
| --- | --- | --- | --- |
| *S. australis*/A12-like | 4 | 432 ± 116 | 1.4 ± 0.2 |
| *S. cristatus* | 11 | 149 ± 68 | 1.0 ± 0.7 |
| *S. gordonii* | 17 | 325 ± 159 | 0.9 ± 0.4 |
| *S. intermedius* | 11 | 417 ± 169 | 0 ± 0 |
| *S. mitis* | 27 | 83.7 ± 125*** | 0.6 ± 0.7 |
| *S. oralis* | 8 | 58.8 ± 37** | 0.28 ± 0.5 |
| *S. oralis* subsp. *dentisani* | 6 | 156 ± 291 | 0.91 ± 0.9 |
| *S. oralis* + *S. oralis* subsp. *dentisani* | 14 | 100 ± 199** | 0.56 ± 0.8 |
| *S. parasanguinis* | 1 | 688 | 1.9 |
| *S. salivarius* | 1 | 46.8 | 0 |
| *S. sanguinis* | 25 | 311 ± 220 | 0.22 ± 0.4 |
| *Strep. species* | 2 | 5.72 ± 2.6 | 0.89 ± 0.8 |

All values are ± SD.
\* p < 0.05 vs. *S. gordonii*
\*\* p < 0.01 vs. *S. gordonii*
\*\*\* p < 0.001 vs. *S. gordonii*

Antagonism towards *S. mutans* was variable within each species and not correlated with mean ADS activity. We compared the mean antagonistic activity of each species to *S. gordonii* for consistency with the ADS activity comparisons, and found that *S. intermedius* and *S. sanguinis* had significantly lower antagonism than *S. gordonii* (Figure 1B, Table 1). *Streptococcus gordonii* and *S. australis*/A12-like isolates all exhibited antagonism towards *S. mutans*, while *S. intermedius* was the only species with no isolates that exhibited antagonism, and all other species had isolates with a range of antagonism from none to high (Figure 1B). The single *S. salivarius* isolate had low ADS activity and was not antagonistic towards *S. mutans*, while the single *S. parasanguinis* had very high ADS activity and low antagonism. Given the wide range of phenotypes within all other species represented here, it is not possible to speculate on whether these characteristics are representative of *S. salivarius* and *S. parasanguinis*. However, previous studies from our lab have shown that *S. salivarius* is an abundant producer ammonia via urease enzyme by which they help to maintain the oral biofilm pH homeostasis (10).

### *Distribution of the ADS operon in the genus* Streptococcus

To better understand the distribution of the ADS operon within the genus *Streptococcus* and to correlate the presence of the ADS operon within our oral isolates, we manually searched for the operon in a custom-built database of *Streptococcus* RefSeq genomes, and performed BLAST searches of the operons found manually against the *Streptococcus* database to look for other genomes with the operon. The entire operon *arcABCDT* and *arcRqueA* were identified in *S. constelatus* (70%), *S. cristatus* (100%), *S. gordonii* (96%), *S. intermedius* (75%), *S. parasanguinis* (70%), and *S. sanguinis* (97%), all of which are oral species (Supplemental Table S2). Isolates lacking *arcR* and *queA* were found in *S. mitis* (3.5 %), *S. oralis* (5.2%), *S. oralis* subsp. *dentisani* (54%) (9% of *S. oralis* + all *S. oralis* subspecies), *S. pneumoniae* (94%) and in *S. sp*. oral taxon 058. The remaining species that had ADS operon genes were *S. anginosus* (10%), *S. canis* (100%), *S. dysgalactiae* (100%), *S. merionis* (100%), *S. pyogenes* (78%), and *S. uberis* (93%) (Supplemental Table S2), yet in all cases the operon was not contiguous or complete. In some species the order of the ADS genes had been rearranged, and in others additional genes or transposons had been inserted without disrupting the genes. Therefore, it remains unclear if the operon is functional in these species.

### Arginine deiminase activity does not correlate with genotype

Arginine deiminase activity in *Streptococcus* is governed by the arginine deiminase operon, which includes 5 structural genes *arcA*, *arcB*, *arcC*, *arcD*, and *arcT*, and the regulatory genes *arcR* and *queA*, which are co-transcribed in the opposite direction from the structural genes (20). The global nitrogen regulator *flp* is also involved in regulating expression of the operon (17), and was annotated *ntcA* in our genomes. We identified each of these genes in our isolates to compare phylogenetic relatedness with ADS activity. The annotation of these genes was not consistent, sometimes *arcD* and *arcT* were annotated as “hypothetical protein” and “putative dipeptidase”, yet we confirmed a full, contiguous operon and associated regulatory genes as described in the methods. All eight genes (*ntcA*, *arcA*-*T*, *arcR*, *queA*) were present in all isolates of *S. australis*/A12-like, *S. cristatus*, *S. gordonii*, *S. sanguinis*, and the single *S. parasanguinis* isolate, but were not detected in the *S. salivarius* isolate or the three unidentified species isolates (Supplemental Table S1). Two of the *S. sanguinis* isolates have a 3-gene insertion between *arcC* and *arcD* that includes *ydgI* and *aspC*, and a duplicated *arcC*, yet this does not appear to have impaired their ADS activity (Supplemental Table S1). Nine of eleven *S. cristatus* isolates had all eight genes and the remaining two isolates had none. Very few isolates of *S. mitis* and *S. oralis* had any genes in the operon, and when present *ntcA* and *arcABCDT* were there, but not *arcR* or *queA*. Six of the 21 *S. mitis* isolates (29%), one of the eight *S. oralis* isolates (12%), and five of the 6 *S. oralis* subsp. *dentisani* (83%) isolates had this part of the operon (Supplemental Table S1). The distribution of the ADS operon in our isolates is similar to its distribution in the RefSeq genomes of these species examined above (Supplemental Table S2). Only one of these genes, *arcD*, tested positive for recombination with phi.

We built phylogenies of three versions of the full operon region including all intergenic regions, one *arcABCDT+arcRqueA* (Figure 2A, B), one *nctA+arcABCDT+arcRqueA* (Figure S3A, B), and one *arcABCDT* (Figure S4A, B) to assess the phylogenetic relatedness of the operon and regulatory elements, and to determine whether the ADS activity of each isolate is related to genotype. We then built individual phylogenies for each of the eight genes (Figure S5). The isolates grouped by species in each operon phylogeny, and gene consensus trees showed similar branching patterns (Figures 2C, S3C, S4C). Branching patterns in each phylogeny closely matched those of the *Streptococcus* genus phylogeny (52). Like the core phylogeny though, the *S. mitis*, *S. oralis* and *S. oralis* subsp. *dentisani* isolates are intermixed within their own clade. Heat maps presenting the mean ADS activity for each isolate aligned with the phylogenies (Figures 2, S3, S4, S5) do not show clear correlations between ADS activity and the species groups or the branching patterns within each species. In the *arcR* phylogeny (Figure S5, S6A), the *S. cristatus* isolates split into two groups because the five isolates in the clade more distant to the *S. gordonii*, *S. intermedius*, and *S. sanguinis* isolates have a very short *arcR* sequence. The short *arcR* sequences are genuinely short and not an artefact of assembly such as truncation due to being located at the end of a contig, and removing them from the phylogeny does not alter the branching patterns delineating the species clades (Figure S6B).

**Figure 2.**
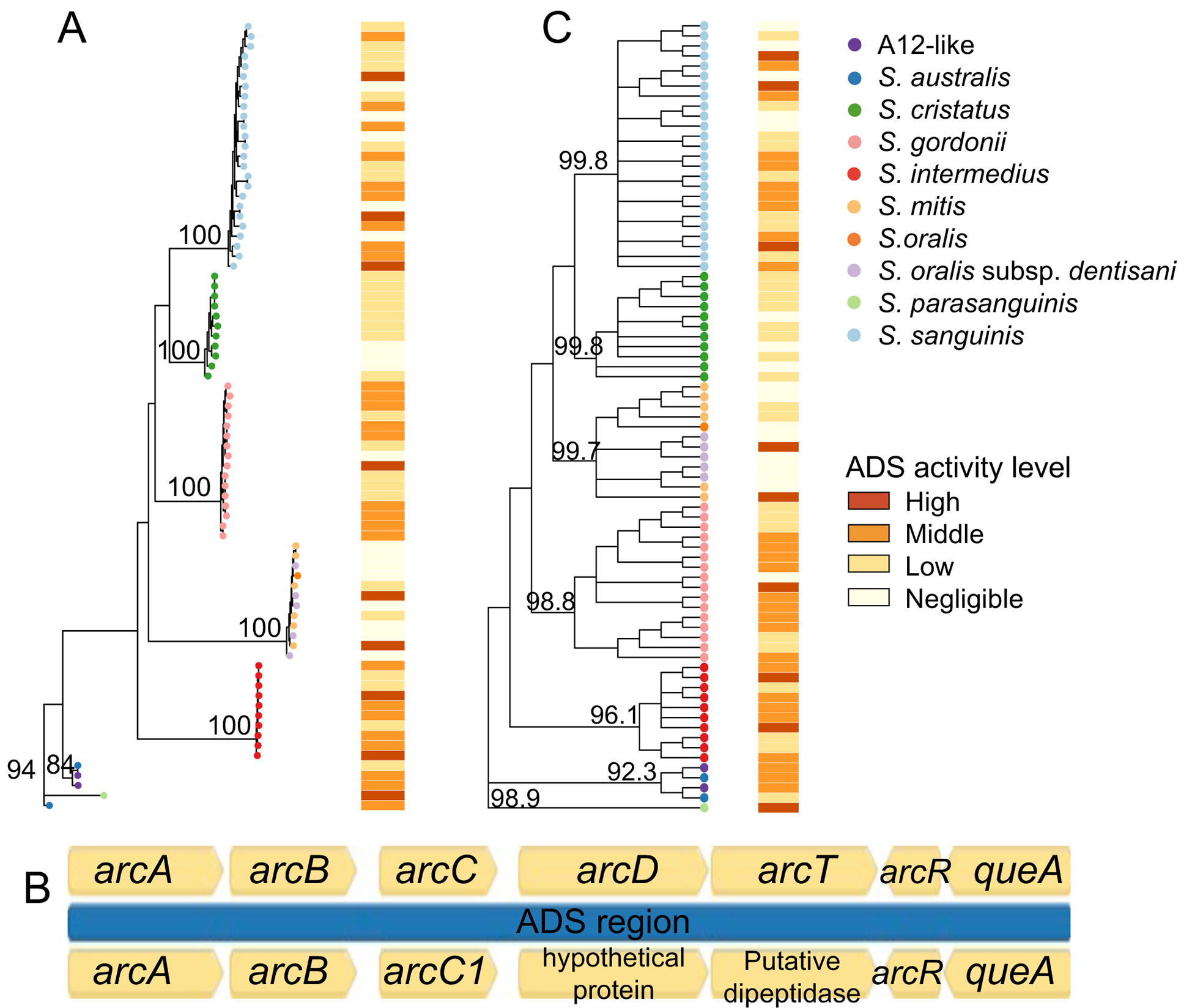
ADS operon genotype and ADS activity level. A. Maximum-likelihood phylogeny of the ADS operon and regulatory elements *arcABCDTRqueA* with heatmap indicating ADS activity level. B. Example of the ADS operon and control elements showing protein-coding and intergenic regions used to build the phylogeny in A, from *S. gordonii* strain Challis (top), and an *S. gordonii* isolate from this study (bottom). Note inconsistencies in gene annotation. C. Gene consensus tree of the individual ADS operon gene trees (*arcA*, *arcB*, *arcC*, *arcD*, *arcT*, *arcR*, *queA*) with heatmap indicating ADS activity level. Bootstrap values (%) are shown on major nodes.

### *Antagonism of* S. mutans *does not correlate with known antagonism-related genotypes*

It was previously shown that targeted loss of the gene for the H_2_O_2_-generating pyruvate oxidase (*spxB*) or the gene for the serine protease challisin of *S. gordonii* DL1 and *Streptococcus* A12, which degrades an intercellular signal molecule for *S. mutans* bacteriocin production, reduces antagonism of these strains towards *S. mutans* (51), so we examined the phylogenetic relatedness of these genes in our isolates. The pyruvate oxidase gene, annotated *pox5* rather than *spxB*, was present in all isolates of *S. australis*/A12-like, *S. cristatus*, *S. gordonii*, *S. oralis*, *S. oralis* subsp. *dentisani*, *S. parasanguinis*, and *S. sanguinis*. We confirmed that this gene is equivalent to *S. gordonii* strain Challisin *spxB* by including that gene in our alignment and building a tree that included *spxB* (Figure S7). All but one *S. mitis* isolate carried the gene and both undefined species isolates carried it, while only a single isolate of *S. intermedius* carried it. The *pox5* phylogeny is not strictly grouped by species like the *arc* gene phylogenies (Figure 3A), and the gene tested positive for recombination with phi. The majority of *S. sanguinis* isolates cluster together, and there is a distinct clade of *S. mitis*/*S. oralis*/*S. oralis* subsp. *dentisani*, yet the remaining isolates form mixed-species clades. The heatmap of mean antagonism activity aligned with the tree in Figure 3A shows no clear relationship between gene phylogeny and antagonistic activity measured in aerobic conditions.

**Figure 3.**
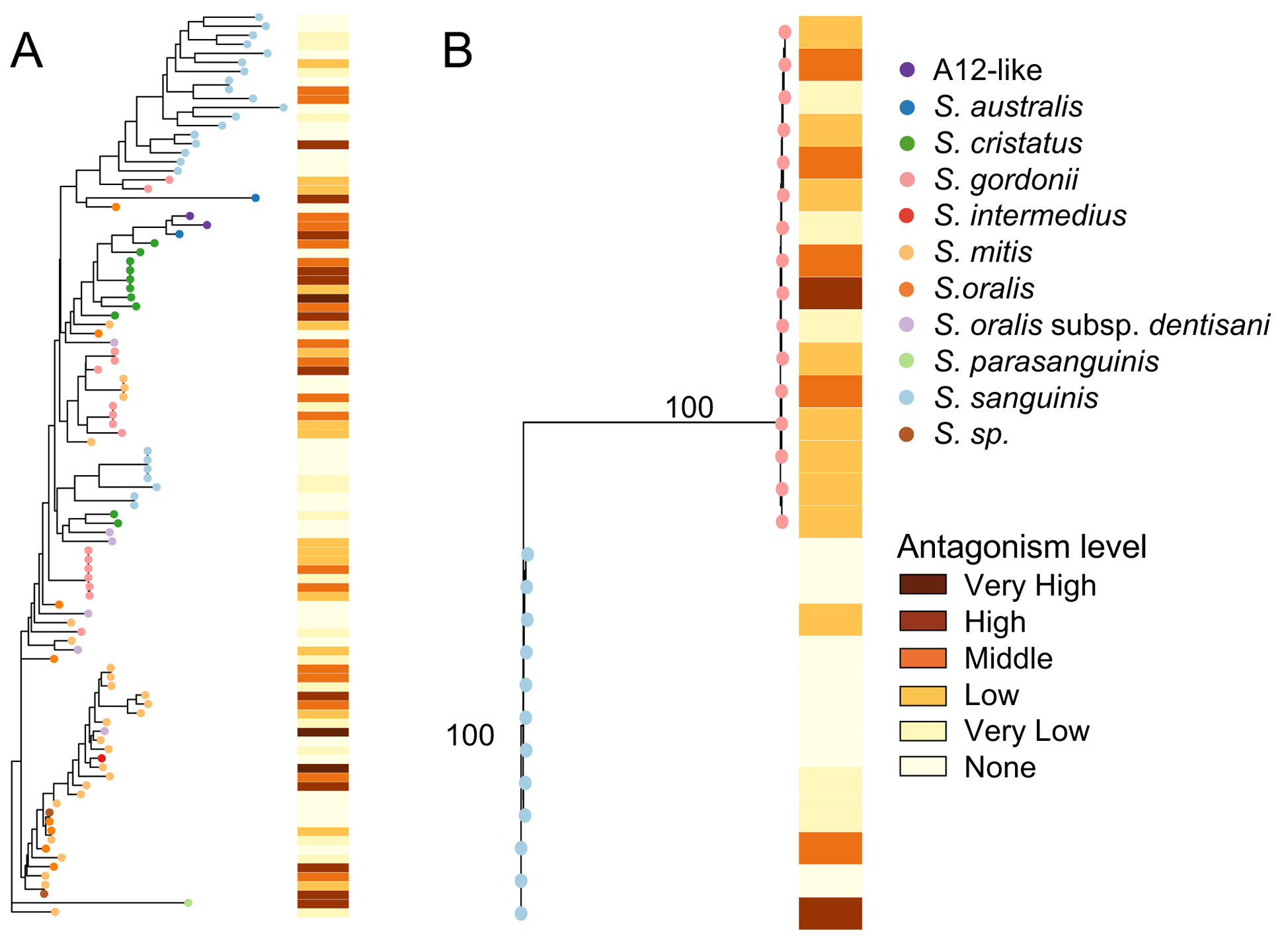
Antagonism-associated genotype and phenotype. A. Maximum likelihood phylogeny of the pyruvate oxidase gene with heatmap indicating level of antagonism towards *S. mutans*. Bootstrap values were <50% for major nodes. B. Maximum likelihood phylogeny of the challisin gene with heatmap indicating level of antagonism towards *S. mutans*. Bootstrap values (%) are shown on major nodes.

The challisin gene was found only in *S. gordonii* and *S. sanguinis* isolates and was annotated *scpA*, a C5a protease. The gene was present in all *S. gordonii* isolates, but only twelve of twenty-five *S. sanguinis* isolates (Figure 3B). In the phylogeny the isolates cluster by species, and the short branches within the species show there is very little variation in the gene sequences. It has not been shown that challisin itself has antagonistic activity, but it might enhance antagonism by diminishing the amount of bacteriocins that *S. mutans* can produce, thereby allowing for better growth and production of antagonistic factors by the commensal. However, although more *S. gordonii* isolates have high antagonism than do *S. sanguinis* isolates, the heatmap of antagonism shows no clear correlation between the challisin phylogeny and antagonism activity. Like *pox5*, *scpA* tested positive for recombination with phi.

### Genus-and species-specific genes potentially involved in ADS activity and antagonism

To search for additional genes that may be involved in ADS activity or *S. mutans* antagonism we screened our isolates using two approaches to detect bacterial *pan-genome-phenotype* - association. We searched for genes associated with these phenotypes across all of our isolates, as well as within each of the species groups. Using the first approach (Scoary) no significant associations between gene clusters and phenotype were found when running 100 permutations and a Benjamini-Hochburg corrected p-value cutoff of 0.05. In contrast, using the second approach (treeWAS) we found sets of genes significantly associated with ADS activity and antagonism across all species, as well as within *S. mitis*, *S. oralis*, and *S. sanguinis* (Supplemental Table S3). Several of the genes associated with ADS activity in all *Streptococcus* isolates we sequenced are involved in arginine processing, including arginine transport system permease *artQ*, arginine decarboxylase, and arginine-binding extracellular protein *artP* precursor (Supplemental Table S3), while several others were involved in outer membrane transport or other seemingly unrelated processes, or were hypothetical proteins. Fewer genes were associated with antagonism in all *Streptococcus* isolates than with ADS activity, and included DNA-binding transcriptional repressor *acrR*, a type-1 restriction enzyme R protein, and a bacteriophage holin.

None of the genes associated with ADS activity or antagonism in the full set of isolates were identified in any of the species-specific tests for association. *S. oralis* and *S. oralis* + *S. oralis* subsp. *dentisani* both had a single gene associated with antagonism, *amiA1* encoding oligopeptide-binding protein AmiA, which was also identified in the *S. mitis* GWAS (Supplemental Table S3). *S. sanguinis* had six genes associated with ADS activity, three of which were hypothetical proteins. One gene, annotated carbamoyl phosphate synthase-like protein, is involved in arginine metabolism, while the relation of the remaining two annotated genes, enterobactin exporter EntS and UDP-N-acetylglucosamine 1-carboxyvinyltransferase 2, to ADS activity is not clear (Supplemental Table S3). A single gene, transcriptional regulator *mtrR1*, was associated with antagonism in *S. sanguinis*. None of the remaining species groups had any genes significantly associated with ADS activity or antagonism.

## Discussion

We performed a genome-wide study of a phylogenetically and phenotypically diverse set of oral streptococci isolated from health-associated supragingival dental plaque to characterize the genotypic basis of variation in ADS activity and antagonism of *S. mutans*. We demonstrated that these two phenotypes vary substantially within and between species, yet the phylogenetic relationship of the genes associated with these phenotypes through earlier studies do not reflect the actual phenotypes. Our results support the observation (18) that differences in transcriptional or translational control may influence the expression of genes responsible for these phenotypes more than the gene sequences themselves.

The ADS operon genes are widely-distributed in *Streptococcus*, but appear to be maintained as a contiguous (and presumably functional) operon predominantly in the oral *Streptococcus* species. This may be directly related to their lifestyle in oral biofilms, which are frequently acidified by other biofilm species, in contrast to other *Streptococus* species such as *S. pyogenes* or *S. uberis*, which are not known to inhabit dense biofilms that are commonly subjected to frequent acidification. However, the operon is clearly functional in the *Streptococcus* species we screened when not contiguous or when lacking regulatory genes. Insertion of 3 genes between *arcC* and *arcD* in two *S. sanguinis* strains was not associated with diminished ADS activity, and despite the *S. mitis* or *S. oralis* strains with the operon missing *arcR* and *queA*, AD activity is still expressed. While the average ADS activities for *S. mitis* and *S. oralis* are lower than other strains, the lack of *arcR* and *queA* does not necessarily explain this, as several strains in *S. gordonii*, *S. cristatus*, and *S. salivarius* have these regulatory genes yet have low ADS activity.

Regulation of the ADS operon is complex and it is not surprising that there is no clear relationship between operon genotype and ADS activity. Expression can be repressed by oxygen, enhanced at low pH, and increased by arginine concentration, and involves several regulatory genes, carbohydrate catabolite repression, and two-component systems (20). Many genes are involved in ADS pathway activation, and this network of regulation may determine expression levels that are unrelated to the sequence of the structural genes. This regulatory network can be identified by functional studies, but not genomic studies alone. In addition, post-transcriptional control of the ADS operon may be important in determining expression levels (20), which again cannot be captured by genomic surveys.

The phylogenetic relationships of antagonism-associated genes pyruvate oxidase, which produces H_2_O_2_ that inhibits *S. mutans* directly, and challisin, which interferes with *S. mutans* bacteriocin production potentially reducing fitness of *S. mutans*, within the isolates that we sequenced do not correlate with the antagonism phenotypes of each isolate, just as we saw for ADS activity genotype and phenotype. There are some clusters of species within the *pox5* pyruvate oxidase phylogeny, but the species groups are much more mixed than was seen with any of the ADS operon genes, which suggests that this gene may be subject to horizontal transfer. The *pox5* gene tested positive for recombination with phi, which supports horizontal transfer between *Streptococcus* species. Similar to the ADS operon gene phylogenies, there is no clear correlation between *pox5* genotype and antagonism phenotype, with the exception of *S. intermedius*. None of the *S. intermedius* isolates were antagonistic towards *S. mutans*, and only a single isolate had the *pox5* gene. Although the challisin gene shows a distinct species-related phylogenetic signal, it shows no correlation with antagonism phenotype. The indistinct relationships between pyruvate oxidase genotype and antagonism as well as challisin genotype and antagonism may again be related to the transcriptional, translation, and/or post-translational control of these genes, or in the case of challisin to differences in substrate specificity of the enzyme.

Our genome-wide association studies did not report associations between ADS activity or antagonism and the genes involved in these phenotypes for which we built phylogenies. Given the complex network regulating ADS operon expression discussed above, this is not surprising. However, several genes that were identified by treeWAS as significantly associated with ADS activity are involved in arginine processing, and therefore the genes identified by this method should be investigated by functional studies for their role in arginine processing and ammonia production. Our small sample size, especially for the individual species groups, may prevent us from finding significantly associated genes, and these GWAS studies should be performed with more isolates to obtain better power, particularly if functional interrelationships can be established with the gene products we identified using TreeWas and ADS levels.

A single isolate each of *S. salivarius* and *S. parasanguinis*, but also included two isolates similar to the recently-described strain A12 (51) based on 16S rRNA gene similarity, were included in our analysis. *S. salivarius* is the most distantly related of the *Strep*. species we included (52), and the isolate we sequenced did not contain any ADS operon genes. None of the 44 RefSeq *S. salivarius* genomes we screened had the ADS operon, so this species may rely instead on the urease gene cluster to produce ammonia to counter drops in pH (10). However, the full urease operon (53) was only present in our *S. salivarius* isolate but none of our other isolates. Urease activity is higher in plaque from caries-free than caries-active adults (27), so this pathway may desirable in probiotic strains. More S*. salivarius* strains will need to be characterized for ammonia production and *S. mutans* antagonism to understand the range of ammonia production in this species, and its potential as a probiotic. In contrast, the *S. parasanguinis* isolate had high ADS activity and moderately antagonized *S. mutans*, and the range of activity in this species should also be further investigated.

*S. australis* and the A12-like isolates, which are phylogenetically closely related (51), have moderate to high ADS activity and *S. mutans* antagonism. This finding supports earlier conclusions that this species may make an excellent probiotic candidate (51). The A12-like isolates are rare, and our plaque screens identified only 2, both of which we included in this study. What the infrequent isolation of A12-like organisms means for the ecology of this organism in the mouth and plaque biofilm is uncertain, and the ability of this organism to integrate and be maintained in the oral biofilm of patients who do not naturally carry it will need to be studied. Unfortunately, a retrospective examination of microbiome studies that used 16S rRNA gene sequencing would not be informative as the 16S rRNA gene of A12-like organisms, *S. australis*, and *S. parasanguinis* share 99% identity with the cannot easily be distinguished. Whether A12-like isolates are strains of *S. australis* or a distinct species is unclear from our core and gene phylogenies. We are in the process of obtaining, characterizing and sequencing more A12-like isolates to clarify the relationship between this species and *S. australis*, and its placement in the phylogeny of the genus *Streptococcus*.

In sum, we have shown that the extensive variation in ADS activity and *S. mutans* antagonism within oral *Streptococcus* species cannot be solely explained by genotypic variation. Complex regulation of these phenotypes may explain the differences within and between species, but cannot be assessed by gene sequence analysis or genome-wide surveys. To develop probiotics that take advantage of ammonia production and growth inhibition of *S. mutans*, strains will need to be carefully selected based on laboratory screening and phenotypic characterization, and not on species designation alone.

## Materials and Methods

### Plaque collection and bacterial strain isolation

Supragingival dental plaque was collected from both children (n=29) and adult (n=11) caries free individuals, those having no clinical evidence of present or prior dental caries activity [decayed, missing and filled teeth (DMFT) = 0]. Informed consent was obtained from all participating subjects (parents in case of children) under reviewed and approved protocols by the Institutional Review Board of the University of Florida Health Science Center (approval number IRB201600154 for children’s study and IRB201600297 for adult study). Children and adult individuals were required to refrain from oral hygiene procedures for 8 and 12 hours prior to the collection of dental plaque, respectively. Plaques samples were collected from teeth surfaces using sterile periodontal curetes, then transferred to sterile, chilled microcentrifuge tubes containing 10 mM sodium phosphate buffer (pH 7.0) and stored at −80° C until further analysis. to isolate cultivable oral *Streptococcus* species (27, 30), plaque samples were dispersed by external sonication (FB120, Fisher Scientific, Hampton, NH, USA) for 2 cycles of 15 seconds with 30 seconds cooling on ice in between. 10 μl of the dispersed plaque samples were then serially diluted in 10 mM sodium phosphate buffer (pH 7.0) and 100 μl of the diluted samples (10^−4^ to 10^−7^) were plated on sheep blood agar (Columbia agar base containing 5% v/v of anticoagulated sheep blood, Difco Laboratories, Michigan, USA) and on BHI (Difco Laboratories) agar. Plates were placed in anaerobic jars (BBL GasPak^TM^ Systems, BD Diagnostics, Md, USA) and incubated at 37° C incubator for 48 hours. Colonies of all clinical isolates from both blood agar and BHI agar plates were collected and further sub-cultured on the same media and incubated subsequently in 5% CO_2_ aerobic incubator until pure colonies were obtained.

### Preliminary species identification by 16S rRNA gene sequencing

To select only *Streptococcus* isolates for biochemical characterization, we sequenced the 16S rRNA gene of our clinical isolates to assign each to a species. An optimized polymerase chain reaction using universal primer set (forward: 5’-AGA GTT TGA TCC TGG CTC AG-3’, reverse: 5’-TAC GGG TAC CTT GTT ACG ACT 3’) was used to amplify the full 16S rRNA gene from each clinical isolate (31). PCR products were then cleaned using Qiaquick PCR cleanup kit (Qiagen, Valencia, Calif., USA) and sequenced by Sanger sequencing at the University of Florida Interdisciplinary Center for Biotechnology (UF-ICBR) for primary identification of isolated bacterial species. A putative species designation for each isolate was determined by a nucleotide BLAST search using the online BLAST search engine at NCBI with default parameters against the 16S ribosomal RNA sequences (Bacteria and Archaea) database, and hit with the highest bit score was selected.

### ADS Activity

All isolated clinical oral streptococci (total 114) were tested for their potential to generate citrulline from arginine via arginine deiminase system (ADS) by a protocol previously validated and published by our group (20). Briefly, a single colony of each clinical isolate was inoculated in tryptone-yeast extract (TY) broth containing 25 mM galactose and 10 mM arginine and incubated overnight at 37° C in 5% CO_2_ aerobic incubator. Overnight cultures were then diluted (1: 20) in the same media until exponential phase (OD_600_= 0.5- 0.6). The cells were harvested, washed and resuspended in 10mM Tris-maleate buffer and further permeabilized with toluene-acetone (1:9) for the measurement of ADS activity. The total protein concentration of the cell suspension was also measured by using BCA protein estimation kit (Pierce, Waltham, Mass., USA) with known bovine serum albumin (BSA) as the standard. ADS activity level in the clinical isolates were normalized to protein content and represented as nanomoles of citrulline generated per minute per milligram of protein. *Streptococcus gordonii* DL1 was used as a reference strain for this assay.

### Competition Assay

BHI agar plates were used for competition assays between commensal streptococci and oral pathogen *Streptococcus mutans* UA159. Overnight cultures from single colonies were adjusted to OD_600_ 0.5. A 6 μl of each culture was then spotted adjacent to each other on agar plates commensal first and UA159 24 hours later. All experiments were performed in aerobic conditions. ImageJ software was used to measure the zone of inhibition (in mm) between competing colonies on plate.

### Statistical Analysis

Statistical differences in mean ADS activity and mean *S. mutans* antagonism were calculated by one-way ANOVA using *S. gordonii* as the reference group with Bonferroni multiple test correction in Prism v7.0d. Graphs were generated using Prism v7.0d.

### DNA isolation and Illumina sequencing

For whole genome shotgun sequencing, genomic DNA were isolated from each commensal streptococci using Wizard^®^ Genomic DNA Purification Kit (Promega, Madison, WI, USA) with some modifications. Briefly, 8 ml overnight culture of each isolate was harvested and resuspended in 480 μl of EDTA and appropriate lytic enzymes were added to the cell suspension (100 μl of 10 mg/ml lysozyme and 2 μl of 5U/ μl of mutanolysin). Cells were harvested after an incubation of 1 hour at 37°C. Then 600 μl of Nuclei Lysis Solution (provided by manufacturer) was added to the cell suspension and the samples were incubated at 80° C for 5 minutes. This step was necessary for the breakdown of the cell wall. RNase was added to cell lysate and incubated about an hour at 37° C to inhibit RNA contamination while purifying genomic DNA. To minimize protein impurities, Protein Precipitation Solution (provided by manufacturer) was added to the RNase treated cell lysate and mixed by vigorous vortexing and incubated on ice for 5 minutes. The total cell lysate was then harvested and the supernatant containing the DNA sample was transferred to a fresh tube containing room temperature isopropanol. The supernatant was rotated at room temperature about an hour or until the thread-like strands of DNA formed a visible mass. Finally DNA was purified in nuclease-free water after two washes in 70% ethanol. Total DNA concentration was measured using NanoDrop™ One Microvolume UV-Vis Spectrophotometer (ThermoFisher Scientific, Waltham, MA, USA) and DNA integrity was determined by 260/280 ratio. DNA from each isolate was prepared for Next-Generation whole genome shotgun sequencing using 2-5ng DNA and the Illumina Nextera-XT library preparation and indexing kit. Libraries were built without deviation from the Illumina recommended protocol, but were normalized by hand, and not with the beads provided in the Nextera-XT kit. Libraries were pooled at a final concentration of 2nM, and sequenced on an Illumina MiSeq using the Illumina MiSeq v2 kit with paired-end sequencing and 250-bp reads. Reads were de-multiplexed by the Illumina software and the raw fastq files were further processed for analysis.

### Read processing, assembly, annotation, and gene clustering

Estimated coverage of each genome was calculated by multiplying the number of reads in each raw fastq file by the read length (250 bases) and then dividing by the average number of nucleotides in a Streptococcus genome (2.9Mbp). Coverage ranged from 20X-300X. Reads were quality-trimmed and genomes assembled using the program A5 (32) with default parameters, and assembly quality was assessed with quast v4.6.3 (33). Assembled genomes were annotated with Prokka v 1.11(34) using a *Streptococcus*-specific amino acid gene sequene database. For gene clustering, Prokka-annotated amino acid fasta files for the isolates we sequenced along with the *Streptococcus mutans* files were concatenated into one file, as well as the arginine deiminase genes *arcA*, *arcB*, *arcC*, *arcD*, *arcT*, *arcR* from *Streptococcus gordonii* strain Challis (NCBI accession CP000725.1) for easy identification of these genes during analysis. Homologous genes among all genomes were delineated using the MCL algorithm (35) as implemented in the MCLBLASTLINE pipeline (available at http://micans.org/mcl). The pipeline used Markov clustering (MCL) to assign genes to homologous clusters based on an all-vs-all BLASTX search with DIAMOND v0.8.22.84 (36) between all pairs of protein sequences using an *E* value cut-off of 1e-5. The MCL algorithm was implemented using an inflation parameter of 1.8. Simulations have shown this value to be generally robust to false positives and negatives (37).

### Species identification by core genome phylogeny

For comprehensive identification, a core genome of single-copy genes present in all isolates we sequenced was determined from the MCL clustering. A total of 608 single-copy core gene clusters were identified, and these were aligned using MUSCLE (38) and checked for recombination using PhiPack (39). Genes identified as recombinant by all three tests (phi, NSS, max χ^2^) were removed from the core gene group. The remaining 425 putatively non-recombinant single-copy core gene alignments were concatenated and the concatenated alignment was used to build a core phylogeny using phyML v. 3.0 (40) with the GTR+G substitution model. Bootstrap support was provided by generating 100 bootstrap replicates. The species designations for each isolate were compared between the 16S rRNA gene and core gene phylogeny and several discrepancies were found. All isolates were assigned to a species based on the core gene phylogeny.

### Distribution of the ADS operon in the genus Streptococcus

To determine the distribution of the contiguous ADS operon within the genus Streptococcus and our sequenced isolates, we built a comprehensive Streptococcus custom BLAST database using the software Geneious v7.0 (https://www.geneious.com). The database was built using Genbank files from RefSeq at NCBI and those generated by Prokka for our isolates. Consequently, the database contained assembled and annotated contigs and information regarding gene synteny was available. All RefSeq *Streptococcus* genomes were downloaded from NCBI on 16 April 2018. Fifty genomes of *S. agalactiae*, *S. equi*, *S. pyogenes*, *S. pneumoniae*, *S. suis*, and *Streptococcus* of unidentified species were randomly selected for inclusion in the database, as there are many more entries of these species in NCBI than the other *Strep*. species. Half of the *Streptococcus mutans* genomes (94 of 187) were included in the database, and all of the *S. oralis* (85) and *S. mitis* (57) genomes were included because we were particularly interested in distribution of the ADS operon in oral *Streptococcus*. We used a total of 1083 *Streptococcus* genomes (Supplemental Table S4) to build the database within the software Geneious v7.0. To obtain a BLAST search query sequence of the contiguous operon we used Geneious to manually search for the *arcA* gene within the genome sequence of S. gordonii strain Challis ADS. This procedure located the operon within a genome and allowed extraction of its contiguous nucleotide sequence.

### Identification of genes involved in the arginine deiminase system

Gene clusters representing genes in the ADS operon (*arcA*, *arcB*, *arcC*, *arcD*, *arcT*) were identified by the presence of *S. gordonii* strain Challis ADS pathway genes in those clusters. The sequences were extracted from the Prokka-annotated fasta files of each isolate by locus tags. We confirmed that each gene was part of the ADS operon in each isolate and not a homologous anabolic counterpart by checking that the locus tag for each gene was sequential with the other ADS operon gene locus tags, as well as confirming that the locus tags of individual genes matched those of the full operon sequence. The *arcR* and *queA* genes were confirmed by checking that the locus tags were sequential with and immediately downstream of the ADS operon, while the regulator *flp* was confirmed by checking that the locus tag was sequential with and immediately upstream of the ADS operon.

The full operon and regulatory genes were identified in our sequenced isolates by manually searching for the operon in a randomly selected representative isolate of each species and performing a BLAST search in Geneious as follows: the annotated genome of one isolate of each species in was searched for the *arcA* annotation and the full operon with regulatory genes *flp*, *arcR* and *queA* was selected and extracted. The extracted full operon was used as the query in a BLAST search against all of our sequenced isolates.

### Identification of genes involved in antagonism

Homologous gene clusters representing the serine protease challisin and the pyruvate oxidase *spxB*, were identified by BLAST search using these genes from *S. gordonii* strain Challis as queries against the genomes of all isolates we sequenced. Homologues of *S. gordonii* strain Challis pyruvate oxidase gene *spxB* were annotated *pox5* in our isolates.

### Association of phenotype and genotype using known genes

Each gene cluster (protease challisin and *spxB*) as well as the full ADS operon was aligned using MAFFT (41) in Geneious and a phylogeny generated using phyML with the GTR substitution model and SPR branch swapping. Brach support was generated via 100 bootstrap replicates. Then, a phylogeny based on the consensus of the separate phylogenies for each gene (gene-trees) (*arcA*, *arcB*, *arcC*, *arcD*, *arcT*, *arcR*, and *queAi*) was constructed using the Triple Construction Method as implemented in the program Triplec (42) (10,000 iterations). This procedure is based on the observation that the most probable three-taxon tree consistently matches the species tree (43). The method searches all input trees for the most frequent of the three possible rooted triples for each set of three taxa. Once found, the set of rooted triples are joined to form the consensus tree using the quartet puzzling heuristic (44). The method has been shown to outperform majority-rule and greedy consensus methods (45). All phylogenies were graphed using the R package ggtree (46). In addition, the alignment for each gene cluster was tested for recombination with PhiPack (37).

### Pan-genome-phenotype association

We searched for genes associated with the phenotypes for our isolates using two genome-wide association approaches: Scoary (47) and treeWAS (48). For Scoary, the genomes of each species were clustered independently using Roary (49) and combined with binary coding of the phenotypes. treeWAS was run using both the individual species clustering obtaining from Roary and combined species clustering obtained using MCLblastline. The phenotypes for treeWAS were coded as both binary and continuous (Supplemental Table S1).

## Acknowledgments

The authors would like to thank Dr. Matthew L. Williams and Kyulim Lee for their help in initial isolation of clinical commensals, Pascale Nehme for assisting during ADS assays, and Kaylin Young for assistance with building libraries. This work was supported by the National Institutes of Health National Institute of Dental and Craniofacial Research R01DE25832 to R.A.B.

## Data Availability

All genomes we sequenced for this study are available to download from the NCBI SRA under accession number PRJNA480251.

## Supplemental Tables

**Table S1.** List of isolates sequenced in this study and their biochemical characteristics.

**Table S2.** List of *Streptococcus* RefSeq genomes used in this study with accession numbers.

**Table S3.** treeWAS results.

---

**Figure S1.**
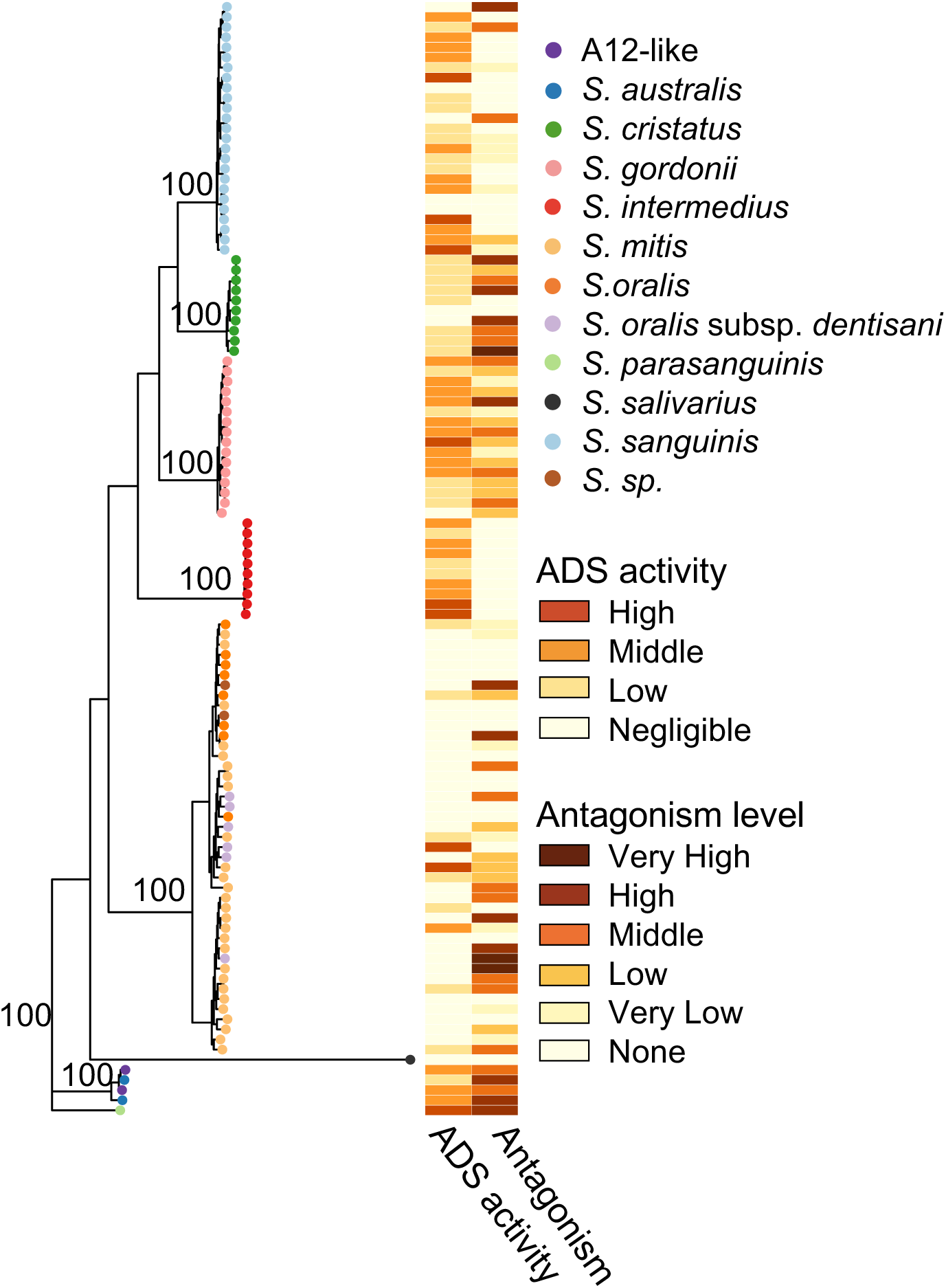
Phylogenetic relationship of isolates sequenced for this study. Maximum likelihood phylogeny based on a core set of 425 putatively non-recombinant genes.

**Figure S2.**
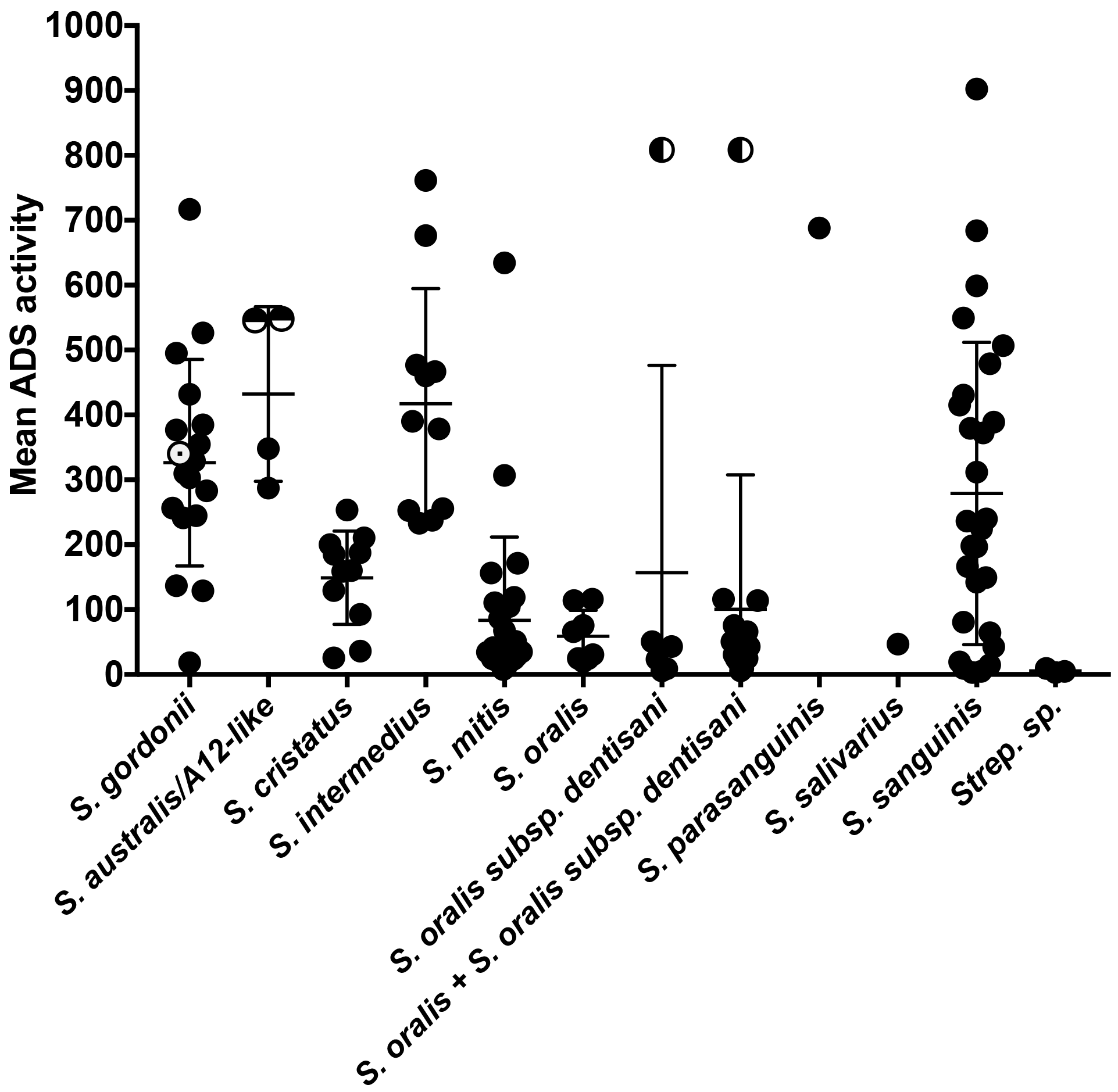
Phenotypic diversity within diverse clinical oral streptococcus isolates. Same as Figure 1A, but with the mean ADS activity of *S. gordonii* DL1 included for reference as a yellow circle with a black dot in its center.

**Figure S3.**
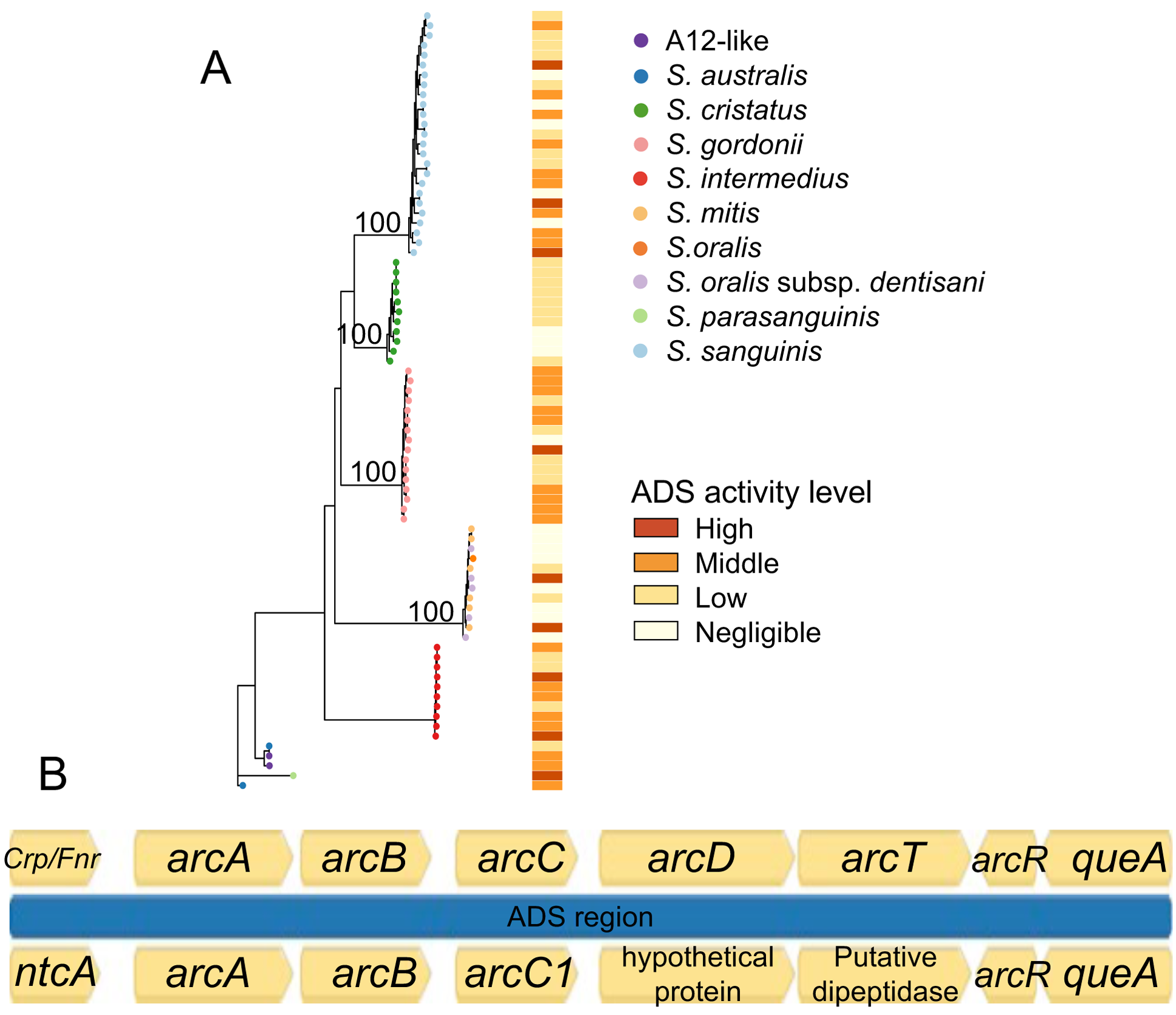
ADS operon and regulatory gene *flp*/*ntcA* genotype and ADS activity level. A. Maximum likelihood phylogeny of the ADS operon *ntcAarcABCDTRqueA* with heatmap indicating ADS activity level. Bootstrap values (%) are shown on major nodes. B. Example of the ADS operon and control elements showing protein-coding and intergenic regions used to build the phylogeny in A, from *S. gordonii* strain Challis (top), and an *S. gordonii* isolate from this study (bottom).

**Figure S4.**
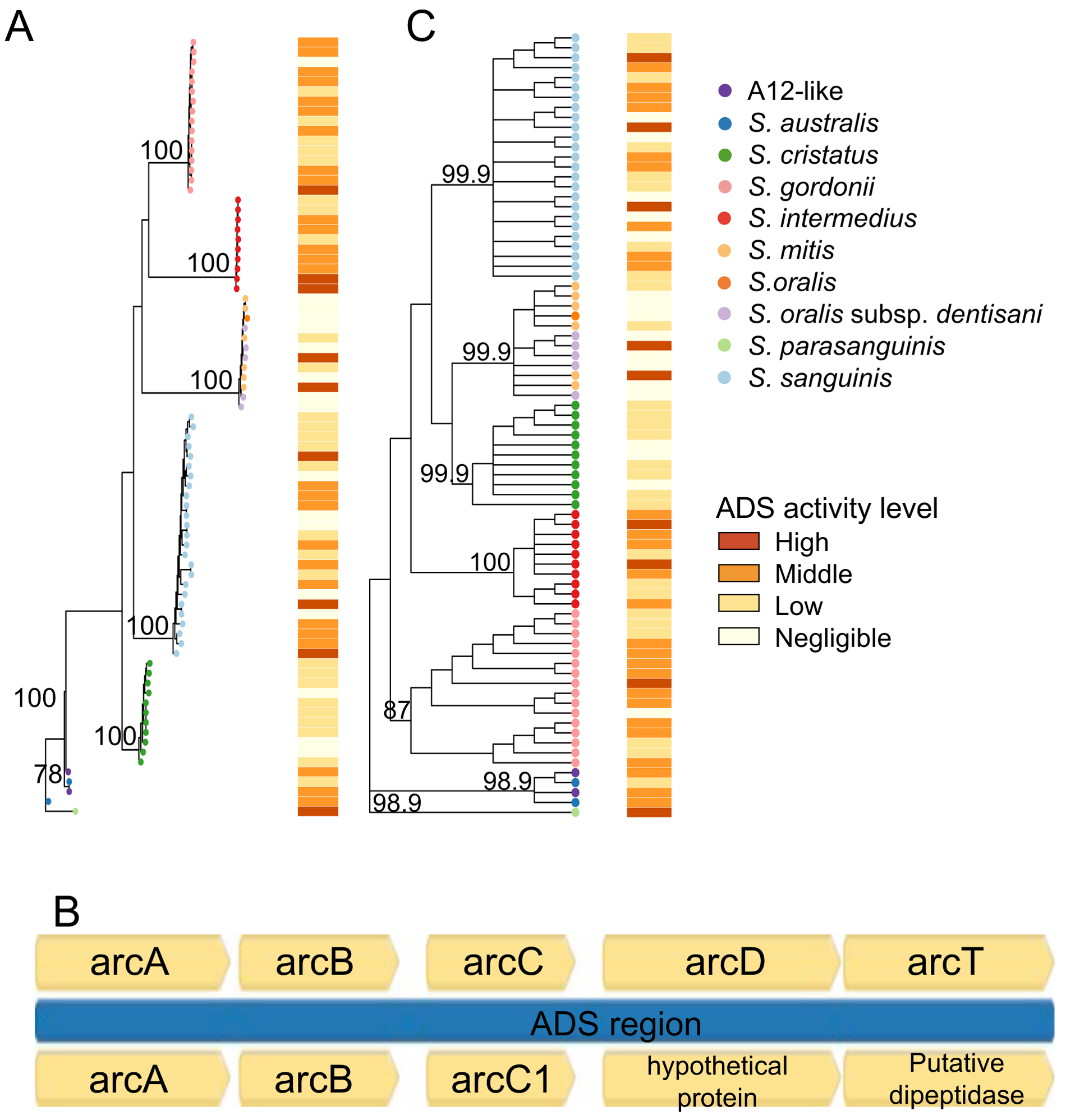
ADS operon genotype and ADS activity level. A. Maximum likelihood phylogeny of the ADS operon *arcABCDT* with heatmap indicating ADS activity level. B. Example of the ADS operon showing protein-coding and intergenic regions used to build the phylogeny in A, from *S. gordonii* strain Challis (top), and an *S. gordonii* isolate from this study (bottom) C. Gene consensus tree of the individual ADS operon gene trees (*arcA*, *arcB*, *arcC*, *arcD*, *arcT*) with heatmap indicating ADS activity level. Bootstrap values (%) are shown on major nodes.

**Figure S5.**
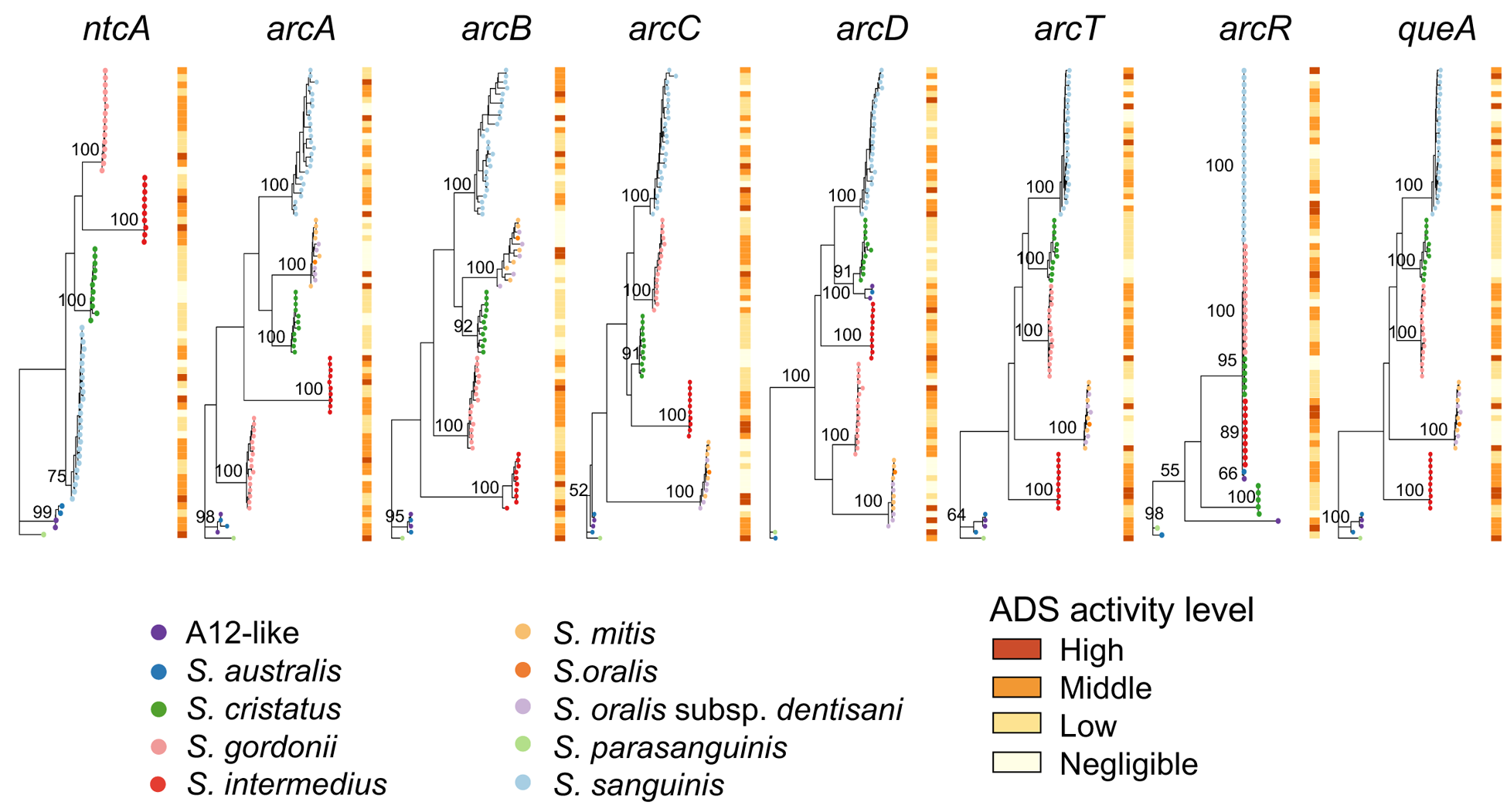
ADS operon gene phylogenies and ADS activity phenotype. Each individual maximum-likelihood gene phylogeny (*ntcA*, *arcA*, *arcB*, *arcC*, *arcD*, *arcT*, *arcR*, *queA*) is presented adjacent to a heat map indicating the ADS activity level of each isolate.

**Figure S6.**
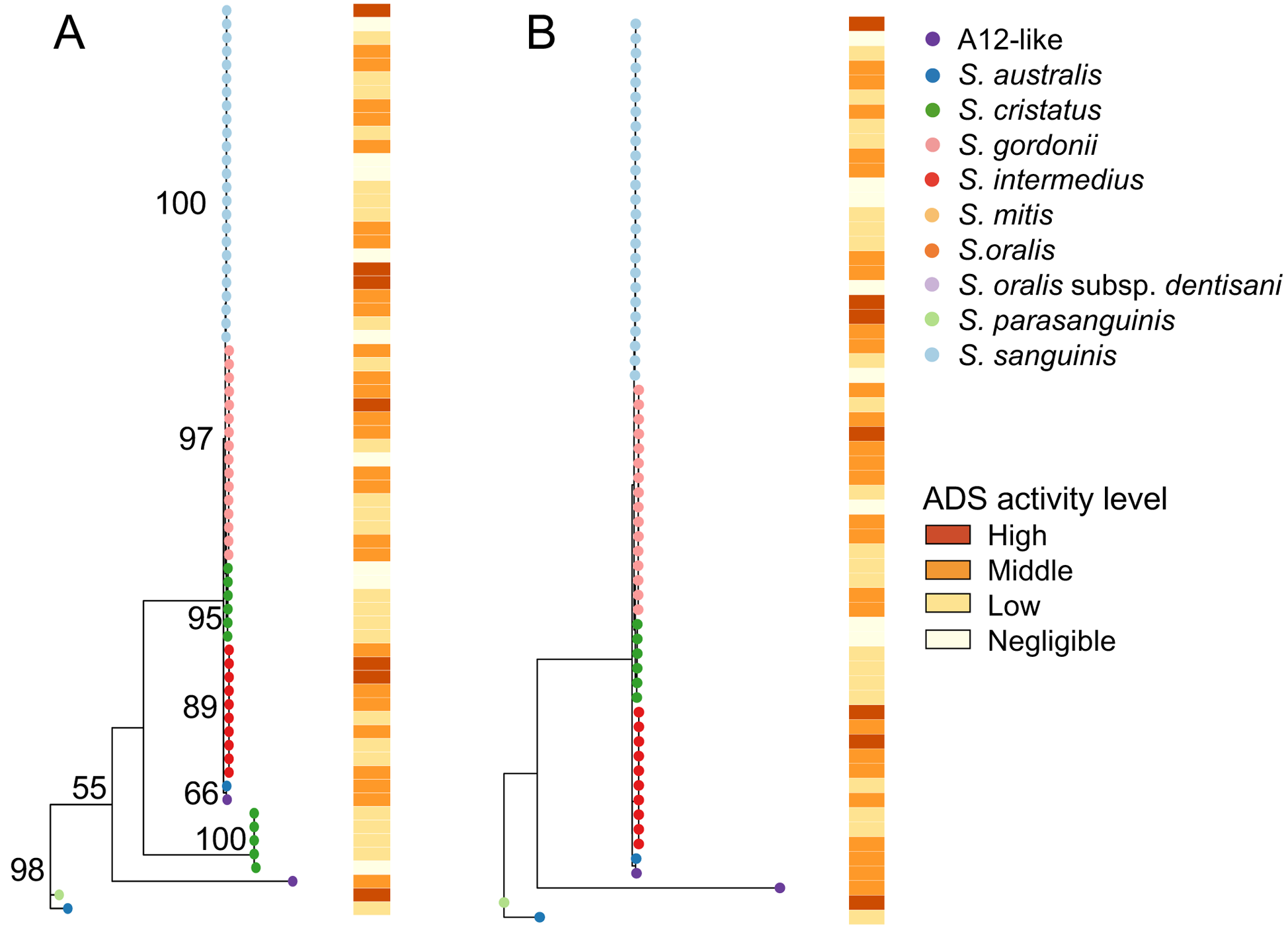
*arcR* gene phylogenies and ADS activity phenotype. A. *arcR* phylogeny for all isolates sequenced in this study with a heat map indicating the ADS activity level of each isolate (same as in Figure S4) B. *arcR* phylogeny for all excluding the 5 *S. cristatus* isolates with short *arcR* sequences, with a heat map indicating the ADS activity level of each isolate.

**Figure S7.**
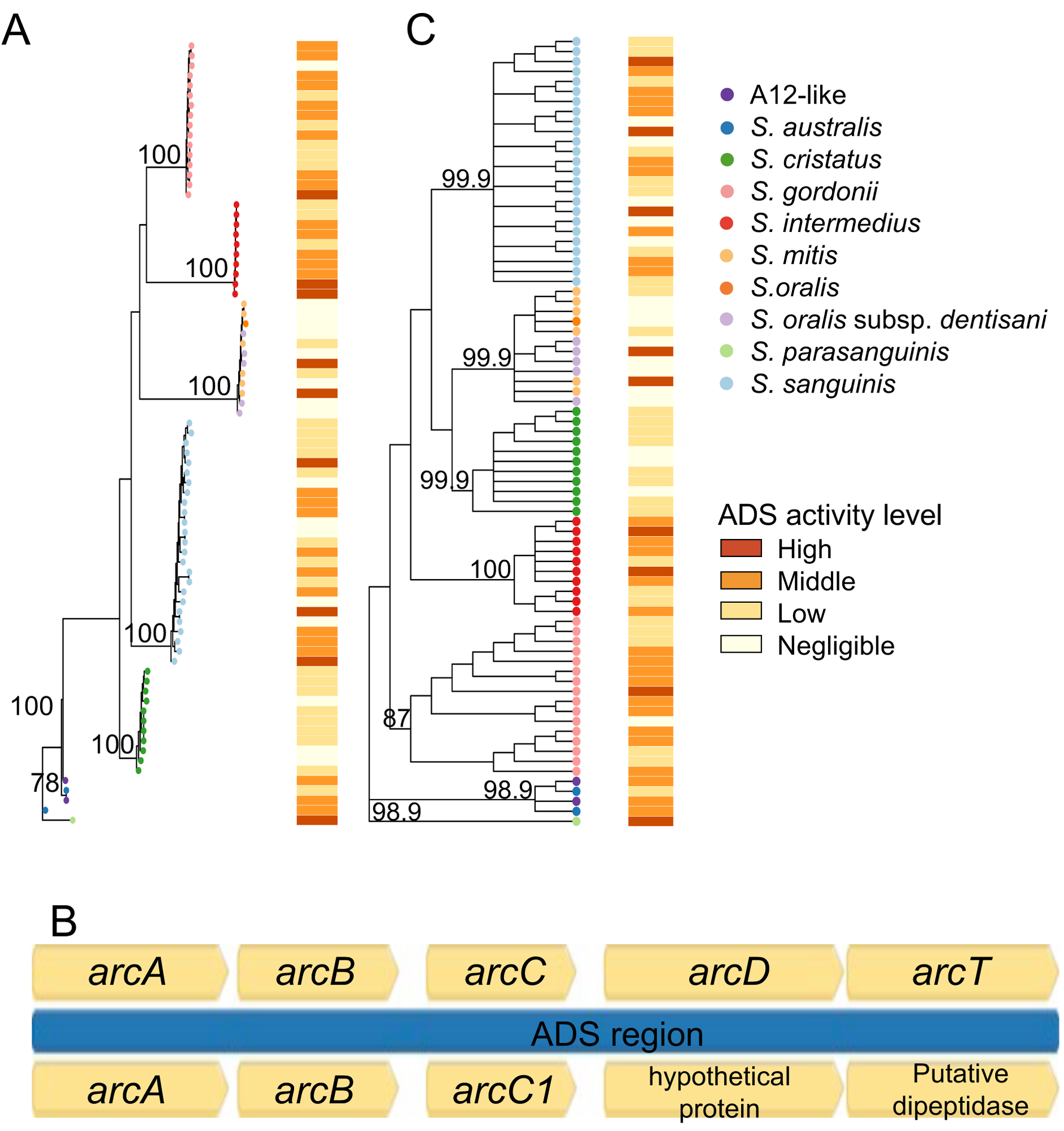
Pyruvate oxidase maximum-likelihood phylogeny with antagonism heat map, same as Figure 3A, but with the *S. gordonii* strain Challis pyruvate oxidase gene *spxB* gene included for reference as a black asterisk (15th from top) for a reference. Bootstrap values were <50% for major nodes.

